# Gene Editing Reveals Obligate and Modulatory Components of the CO_2_ Receptor Complex in the Malaria Vector Mosquito, *Anopheles coluzzii*

**DOI:** 10.1101/2020.05.13.094995

**Authors:** Feng Liu, Zi Ye, Adam Baker, Huahua Sun, Laurence J. Zwiebel

**Author notes:** denotes equal contributions (Feng Liu, Zi Ye).

## Abstract

The sensitivity to volatile carbon dioxide (CO_2_) produced by humans and other animals is a critical component in the host preference behaviors of the malaria vector mosquito *Anopheles coluzzii*. The molecular receptors responsible for the ability to sense CO_2_ are encoded by three putative gustatory receptor (*Gr*) genes (*Gr22,23,24*) which are expressed in a distinctive array of sensory neurons housed in maxillary palp capitate peg sensilla of *An. coluzzii*. Despite the identification of these components and subsequent studies, there is a paucity of understanding regarding the respective roles of these three GRs in the mosquito’s CO_2_ transduction process. To address this, we have used CRISPR/Cas9-based gene editing techniques combined with *in vivo* electrophysiological recordings to directly examine the role of *Gr22,23,24* in detecting CO_2_ in *An. coluzzii*. These studies reveal that both *Gr23* and *Gr24* are absolutely required to maintain *in vivo* CO_2_ sensitivity while, in contrast, *Gr22* knock out mutants are still able to respond to CO_2_ stimuli albeit with significantly weaker sensitivity. Our data supports a model in which *Gr22* plays a modulatory role to enhance the functionality of *Gr23/24* complexes that are responsible for CO_2_ sensitivity of mosquitoes.

## Introduction

The malaria parasite *Plasmodium falciparum* infected an estimated 228 million people worldwide in 2018, accounting for more than 400,000 deaths (WHO, 2019). Accordingly, controlling *Anopheles* mosquitoes, which are the sole vectors for transmitting human malaria, has long been a formidable concern to public health. Blood feeding is a requisite activity for the reproductive cycle in female *Anopheles* mosquitoes and in that context serves as the mode of transport for *P. falciparium* to transit between mosquito vectors and host animals (Aly et al., 2009; Beier, 1998; Whitten et al., 2006).

The mosquito host-seeking process relies heavily on a broad array of sensory cues released from the human body including heat, natural odors, and CO_2_ which in particular has proven to play a critical role in long-range host seeking by mosquitoes (Cooperband & Carde, 2006; Healey & Copland, 1995; Lorenz et al., 2013). This is exemplified by the ability of mosquitoes to detect CO_2_ at distances of nearly 50 meters for *An. arabiensis* (Lorenz et al., 2013) and 150 centimeters for Culex mosquitoes (Cooperband & Carde, 2006). Numerous studies have shown that when CO_2_ is combined with other sensory cues such as heat or odors in creating mosquito traps, the trapping efficiency is considerably higher than when using the heat or odor alone (Geier & Boeckh, 1999; Kline et al., 2012; Spitzen et al., 2008). Indeed, CO_2_ is the most common component of odor baits found in commercial mosquito traps, which illustrates the mosquito’s remarkable attraction to it in the field (Hoel et al., 2007; Kline et al., 1990; Roiz et al., 2012).

Mosquito CO_2_ detection is primarily mediated through the maxillary palp which is an important accessory olfactory head appendage (Grant et al., 1995; Lu et al., 2007; Syed & Leal, 2007). While there appears to be only a single type of olfactory sensillum on the maxillary palp, the capitate-peg (cp), recent studies suggest there may be a degree of functional heterogeneity among these seemingly uniform sensory structures (Ye et al., 2020). That said, there are three different neurons housed in the cp sensillum only one of which, the cpA neuron, is responsible for detecting CO_2_ and other cues; the two other neurons (cpB/C) respond to a range of odors associated with human emanations and other biological sources (Lu et al., 2007). Molecular studies in *Anopheles gambiae* revealed three gustatory receptor genes, *Grs 22, 23*, and *24*, are expressed in the cpA neuron and play a key role in its firing in response to CO_2_ stimulation (Lu et al., 2007).

Phylogenetic analyses reveal the three CO_2_-sensitive *Gr* genes are highly conserved across insect taxa, extending across dipterans and broad evolutionary distances to include Coleoptera, Hemiptera, Hymenoptera and Lepidoptera (Benoit et al., 2016; McMeniman et al., 2014; Robertson & Kent, 2009; Terrapon et al., 2014). Paradoxically, while the genomes of most insect species contain homologs of all three CO_2_-sensitive *Gr* genes, the common fruit fly (*Drosophila melanogaster*) has only two CO_2_-sensitive *Gr* genes, *Gr21a* and *Gr63a* which are orthologous to *Gr1/22* and *Gr3/24* in *Aedes* and *Anopheles* mosquitoes, respectively (Kwon et al., 2007). Considering the evolutionary advancement of *Drosophila* in Hexapods this is likely to be the result of a selective gene-loss event for the third (*Gr2/23*) CO_2_ receptor gene (Misof et al., 2014).

Nevertheless, while *D. melanogaster* only possess two CO_2_-sensitive GRs on the CO_2_-sensitive ab1C neuron, this complex is clearly sufficient for sensing environmental CO_2_ (Jones et al., 2007). Heterologous over-expression of *Gr21a* and *Gr63a* in the *Drosophila* ab3A neuron conferred cryptic CO_2_ sensitivity, further confirming these genes are sufficient in mediating CO_2_ responses in *Drosophila* (Kwon et al., 2007). These results raise questions as to the requirement and role of the third *Gr* gene in the CO_2_-receptor triad complex that is likely to be active in other insect species. These issues are especially salient for mosquitoes which rely heavily on CO_2_-responses for obtaining reproductively essential blood meals and largely forms the basis of their vectorial capacity insofar the transmission of malaria and a range of arboviral diseases.

To answer these questions, we have taken an *in vivo* approach in the Afrotropical malaria vector mosquito *An. coluzzii* using gene-editing techniques to knock out each CO_2_-sensitive *Gr* gene and directly characterize the cpA neuron’s response to CO_2_ in mutant mosquitoes. Comprehensive interrogation of the CO_2_-sensitive neurons of *Gr22, 23*, and *24* mutant mosquitoes reveals that Gr22 plays a modulatory role to enhance the essential and irreplaceable functionality of Gr23/24 complexes that together are responsible for CO_2_ sensitivity of *An. coluzzii*. Our *in vivo* studies align with similar models derived from insect and *Xenopus* heterologous expression studies of the three CO_2_-sensitive Grs from *Aedes aegypti* and *Culex quinquefaciatus* (Kumar et al., 2020; Xu et al., 2020), respectively, suggesting that mosquitoes utilize a similarly distinctive mechanism for CO_2_ reception.

## Material and Methods

### Mosquito maintenance

*Anopheles coluzzii* (SUA 2La/2La), previously known as *Anopheles gambiae sensu stricto* “M-form”(Coetzee et al., 2013), originated from Suakoko, Liberia, were reared as described (Fox et al., 2001; Qiu et al., 2004) and 5-to 7-day-old females that had not been blood fed were used for all experiments. All mosquito lines were reared at 27°C, 75% relative humidity under a 12:12 light-dark cycle and supplied with 10% sugar water in the Vanderbilt University Insectary.

### Electrophysiology

Single sensillum recordings (SSR) were carried out as previously described (Liu et al., 2013) with minor modifications. Mated female mosquitoes (4-10 days after eclosion) were mounted on a microscope slide (76 × 26 mm) (Ghaninia et al., 2007). Maxillary palps were fixed using double-sided tape to a cover slip resting on a small bead of dental wax to facilitate manipulation and the cover slip was placed at approximately 30 degrees to the mosquito head. Once mounted, the specimen was placed under an Olympus BX51WI microscope and antennae were viewed at high magnification (1000×). Two tungsten microelectrodes were sharpened in 10% KNO2 at 10 V. The grounded reference electrode was inserted into the compound eye of the mosquito using a WPI micromanipulator and the recording electrode was connected to the pre-amplifier (Syntech universal AC/DC 10x, Syntech, Hilversum, The Netherlands) and inserted into the shaft of the olfactory sensillum to complete the electrical circuit to extracellularly record OSN potentials (Den Otter et al., 1980). Controlled manipulation of the recording electrode was performed using a Burleigh micromanipulator (Model PCS6000). The preamplifier was connected to an analog-to-digital signal converter (IDAC-4, Syntech, Hilversum, The Netherlands), which in turn was connected to a PC-computer for signal recording and visualization.

Customized CO_2_ tanks at different concentrations (0.001%, 0.005%, 0.01%, 0.05%, 0.1%. 0.5%, 1%) were purchased from Airgas Inc. (Nashville, TN). Compounds with highest purity, typically ≧99% (Sigma-Aldrich) were diluted in dimethyl disulfide (DMSO) to make 1% v/v (for liquids) or m/v (for solids) solutions. For each compound, a 10μL portion was dispersed onto a filter paper (3 × 10mm) which was then inserted into a Pasteur pipette to create the stimulus cartridge. A sample containing the solvent alone served as the control. The airflow across the antennae was maintained at a constant 20 mL/s throughout the experiment. Purified and humidified air was delivered to the preparation through a glass tube (10-mm inner diameter) perforated by a small hole 10cm away from the end of the tube into which the tip of the Pasteur pipette could be inserted. The stimulus was delivered to the sensilla by inserting the tip of the stimulus cartridge into this hole and diverting a portion of the air stream (0.5 L/min) to flow through the stimulus cartridge for 500ms using a Syntech stimulus controller CS-55 (Syntech, Hilversum, The Netherlands). The distance between the end of the glass tube and the antennae was ⩽1cm. The CO_2_ stimulus was pulsed through a separate delivery system that delivered controlled pulses using a PSM 8000 microinjector (WPI, Sarasota, FL., variable 5 mL s^−1^) into the same humidified airstream, from CO_2_ tanks of different concentrations.

Signals were recorded for 10s starting 1 second before stimulation, and the action potentials were counted off-line over a 500ms period before and after stimulation. Spike rates observed during the 500ms stimulation were subtracted from the spontaneous activities observed in the preceding 500ms and counts recorded in units of spikes/s.

### CRISPR-Cas9 gene editing

The CRISPR gene targeting vector was a generous gift from the lab of Dr. Andrea Crisanti of Imperial College London, UK (Hammond et al., 2016). The single guide RNA (sgRNA) sequences for each CO_2_ receptor gene were designed for high efficiency using the Chopchop online tool (http://chopchop.cbu.uib.no/), commercially synthesized (Integrated DNA Technologies, Coralville, IA) and subcloned into the CRISPR vector via Golden Gate cloning (New England Biolabs, Ipswich, MA). The homology templates for *Gr23* and *Gr24* were constructed based on a pHD-DsRed vector (a gift from Kate O’Connor-Giles; Addgene plasmid #51434; http://n2t.net/addgene:51434; RRID:Addgene 51434). Here, the 2kb homology arms extending in either direction from the double-stranded break (DSB) sites were PCR amplified and sequentially inserted into the AarI/SapI restriction sites on the vector. The DsRed sequence was replaced by the ECFP fluorescence marker sequence to construct a pHD-ECFP vector for the knockout of *Gr22*.

The microinjection protocol was carried out as described (Pondeville et al., 2014; Ye et al., 2020). Briefly, newly laid (approximately 1hr-old) embryos of wild type *An. coluzzii* were immediately collected and aligned on a filter paper moistened with 25mM sodium chloride solution. All the embryos were fixed on a coverslip with double-sided tape and a drop of halocarbon oil 27 was applied to cover the embryos. The coverslip was further fixed on a slide under a Zeiss Axiovert 35 microscope with a 40X objective. The microinjection was performed using an Eppendorf FemtoJet 5247 (Eppendorf, Enfield, CT) and quartz needles prepared using a customized protocol (Sutter Instrument, Novato, CA). The gene targeting vector and the homology template were diluted to 300ng/μL each and co-injected to maximum capacity/embryo. Injected embryos were subsequently placed in deionized water with artificial sea salt (0.3g/L) and thereafter reared under normal VU insectary conditions.

The first generation (G0) of injected adults were separated based on gender and crossed to 5X wild type gender-counterparts. Their offspring (F1) were manually screened for DsRed/ECFP-derived red/cyan eye fluorescence using an Olympus BX60 Compound Fluorescent Microscope (Olympus, PA). Red/cyan-eyed F1 males were individually crossed to 5X wild type females to establish a stable mutant line. DNA extraction was performed using Qiagen Gel Extraction protocols (Qiagen, Germantown, MD) and genomic DNA templates for PCR analyses of all individuals were performed (after mating) to validate the fluorescence marker insertion using primers that cover DSB sites (Table S1). Salient PCR products were sequenced to confirm the accuracy of the genomic insertion. Heterozygous mutant lines were thereafter back-crossed to wild type *An. coluzzii* for at least 3 generations before putative homozygous individuals were manually screened for DsRed/ECFP-derived red/cyan eye fluorescence intensity.

Putative homozygous mutant individuals were mated to each other before being sacrificed for genomic DNA extraction and PCR analyses (as above) to confirm their homozygosity.

## Results

### Generation of mosquito mutant lines

In order to generate loss of function mutant mosquito lines for each of the Gr22, 23,24 CO_2_ receptors, guide RNAs (gRNA) were designed to target early exon coding regions to generate truncated and nonfunctional proteins (Figure 1A, B) using the Chopchop online tool (Labun et al., 2019). At the same time, visible markers were inserted into the DSB site of each *Gr* gene target to drive the expression of red eye color in *Gr23* and *Gr24* mutants, and cyan eye color in *Gr22* mutants to facilitate selection (Fig.1C).

**Figure 1.**
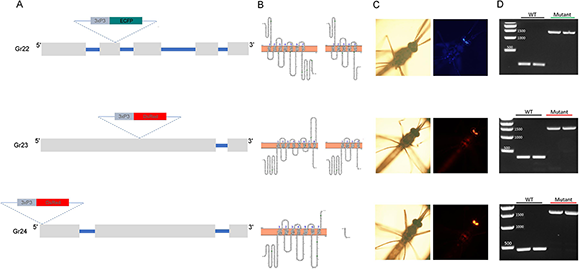
Generation of three *Gr* mutants using CRISPR/Cas9 gene editing. **(A)** Schematic diagram showing the gRNA target site and marker gene (ECFP for *Gr22*, DsRed for *Gr23* and *Gr24*; 3xP3 is the promoter sequence) inserted into the exon of each *Gr* gene. **(B)** Truncation of amino acid sequences after insertion of marker gene in the open reading frame of *Gr* genes. Left: model for secondary structure of normal Gr protein; Right: model for secondary structure of truncated Gr protein. **(C)** Florescence marker presenting in the compound eye of mutant mosquitoes with a specific *Gr* gene knockout. **(D)** Genotyping of the mutant mosquito lines using a pair of primers which amplify a PCR product across the DSB site.

Heterozygous mutant mosquito lines were thereafter back crossed to wild-type mosquitoes for at least three generations before homozygous *Gr* gene mutant lines were established by self-crossing. Each mutant line was molecularly confirmed as homozygotic insertions into each respective *Gr* gene by PCR with a pair of primers to amplify genomic DNA sequences covering the DSB sites of each *Gr* gene, which also served to confirm the specific insertion of eye color markers (Figure 1D). It is noteworthy that while *Gr22*^*-/-*^ and *Gr24*^*-/-*^ mosquitoes appear fully capable of successful breeding leading to the stable generation of laboratory colonies, *Gr23*^*-/-*^ mosquitoes have thus far failed to successfully self-reproduce even after nine generations of backcrosses with wildtype partners.

### Both *Gr23* and *Gr24* are obligatory for CO_2_ detection in the cpA neuron

In light of the absence of a *Gr23* ortholog in *D. melanogaster*, we initially determined the functional relevance of *Gr23* in *An. coluzzii* by assessing neuronal activity in the CO_2_ sensitive maxillary palp cp sensillum (Figure 2A). Here, the cpA neuron in wild-type females display pre-stimulus background activity as well as robust responses to 1% CO_2_ along with a characteristic post-CO_2_ stimulation latency together with low threshold sensitivity to 1-Octen-3-ol (10^−7^ v/v) in cpB/C neurons (Figure 2A). As expected, heterozygous *Gr23*^+/-^ mutants displayed near wild-type responses across several parameters including background activity, CO_2_ sensitivity, and post-response latency (Figure 2B). In contrast, homozygous *Gr23*^−/-^ mutant maxillary palp cpA neurons display no background activity and are completely insensitive to stimulation with 1% CO_2_ even though their cpB/C neurons display wild-type levels of background firing and low threshold sensitivity to 1-Octen-3-ol (Figure 2C, Supplemental Figure S1). These data suggest that, unlike *Drosophila* which lacks a *Gr23* ortholog, in *An. coluzzii*, this gene encodes an essential component for detection of CO_2_ in the cpA neuron and that complexes made up of solely *Gr22*/*Gr24* are insufficient for robust detection of CO_2_ in *Anopheline* mosquitoes.

**Figure 2.**
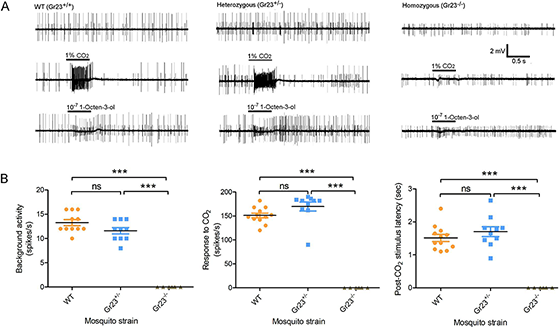
Neuronal responses of wild-type, *Gr23* heterozygous and homozygous mutant mosquitoes when challenged with CO_2_. **(A)** Representative recordings from capitate peg sensilla in the maxillary palp of wild-type, *Gr23* heterozygous and homozygous mutant mosquitoes; Top to bottom: background activity, response to 1% CO_2_ and response to 10^−7^ (v/v in paraffin oil) 1-Octen-3-ol. **(B)** Statistical analysis of background activity (unstimulated spike frequency), firing frequency of cpA neuron in response to CO_2_ challenge, and latency of cpA neuron after CO_2_ challenge. Nonparametric Mann-Whitney test was applied in the statistical analysis with P<0.05 (*), P<0.01 (**) and P<0.001 (***) as significant differences.

*Gr24* homologs are found in the genomes of various insect species and moreover are highly conserved across most, if not all, mosquitoes. Previous studies in *Ae. aegypti* revealed the essential role of *Gr3* (the Anopheles *Gr24* ortholog) in cpA sensitivity to CO_2_ (McMeniman et al., 2014) and, not surprisingly, this phenotype is also seen in our electrophysiological characterization of *An. coluzzii Gr24* mutant females. Here, the cpA neuron in heterozygous *An. coluzzii Gr24*^+/-^ mutants display wild-type background activity as well as robust responses to 1% CO_2_ stimulation (Figure 3A, B) while cpA neurons of homozygous *Gr24*^−/-^ mutants manifest no background activity and are unresponsive when stimulated with 1% CO_2_ (Figure 3A, B). As was the case for *Gr23*, these data confirm that *Gr24/Gr3* is similarly absolutely required for maintaining cpA function and CO_2_ sensitivity in mosquitoes.

**Figure 3.**
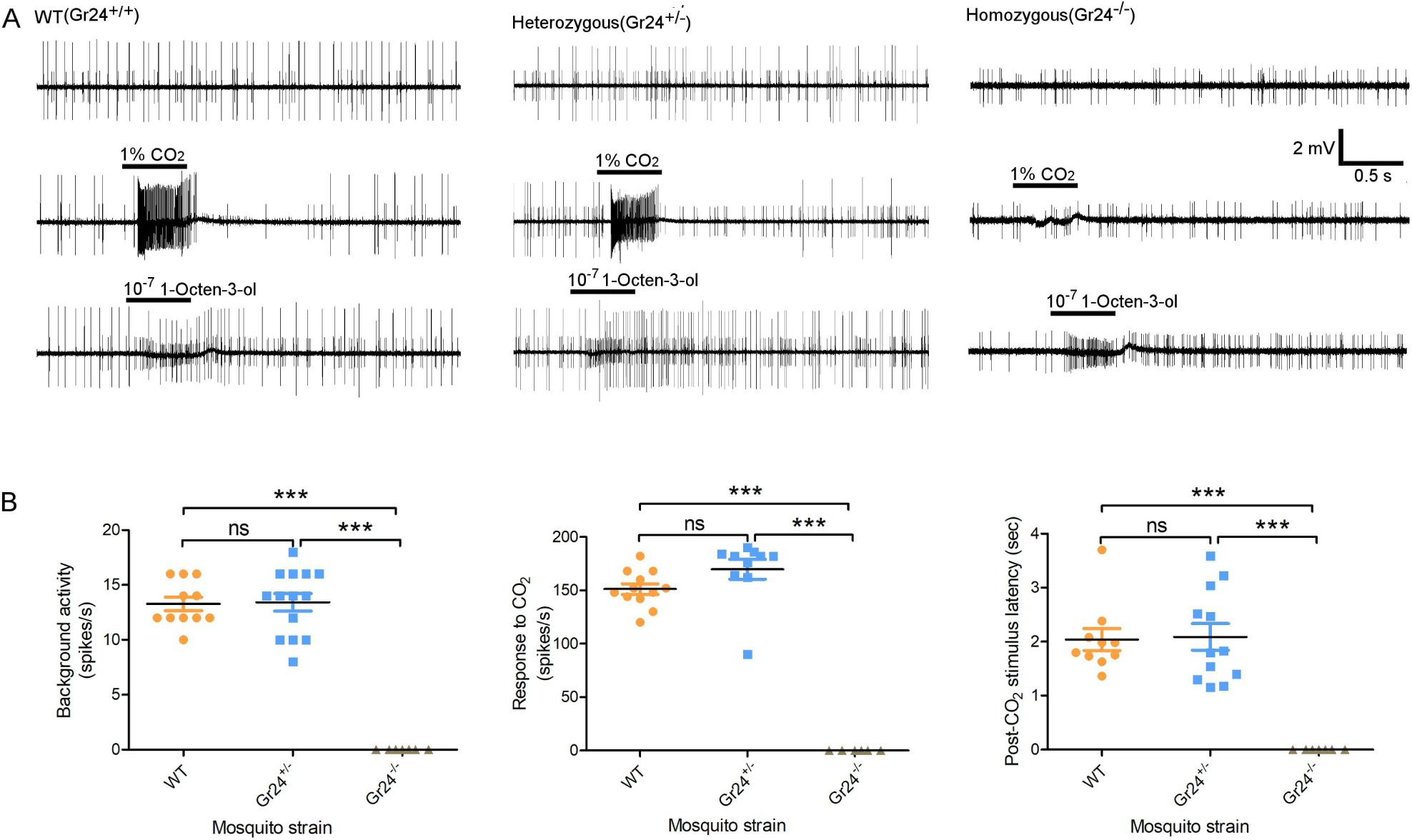
Neuronal responses of wild-type, *Gr24* heterozygous and homozygous mutant mosquitoes when challenged with CO_2_. **(A)** Representative recordings from capitate peg sensilla in the maxillary palp of wild-type, Gr24 heterozygous and homozygous mutant mosquitoes; Top to bottom: background activity, response to 1% CO_2_ and response to 10^−7^ (v/v in paraffin oil) 1-Octen-3-ol. **(B)** Statistical analysis of background activity, firing frequency of cpA neuron in response to CO_2_ challenge, and latency of cpA neuron after CO_2_ challenge. Nonparametric Mann-Whitney test was applied in the statistical analysis with P<0.05 (*), P<0.01 (**) and P<0.001 (***) as significant difference.

### *Gr22* is required for comprehensive CO_2_ detection in the cpA neuron

Recent heterologous expression-based studies have reported somewhat conflicting results on the modulatory role of *Gr22* orthologs in two mosquito species where they are both known as *Gr1* (Kumar et al., 2020; Xu et al., 2020). *Xenopus* oocyte-based studies suggest that in *Cx. quinquefasciatus, Gr1* serves as an inhibitor to fine-tune responses to CO_2_ (Xu et al., 2020). In *Drosophila* empty neuron-based studies, *Ae. aegypti Gr1* acts as both an activator of CO_2_ detection and an inhibitor of sensitivity to pyridine and potentially other non-CO_2_ odors (Kumar et al., 2020). Cognizant of the need to examine these questions in *An. coluzzii* as well as the potential limitations of studies using *in vitro*/heterologous expression systems, we elected to work *in vivo* to directly examine the role of Gr22 in CO_2_ sensitivity of peripheral neurons. As expected, maxillary palp cpSSRs of heterozygous *Gr22*^*+/-*^ mutants showed no significant differences when compared to wild-type preparations insofar as their background activity or in their responses to CO_2_ or 1-Octen-3-ol (Figure 4). However, in contrast to its near absolute silence in *Gr23*^*-/-*^ and *Gr24*^*-/-*^ mutants, the cpA neuron in *Gr22*^*-/-*^ mutants displays modest, albeit significantly reduced background activity, sensitivity to CO_2_ and post-stimulus latency when compared to both wild-type and heterozygous mosquitoes (Figure 4).

**Figure 4.**
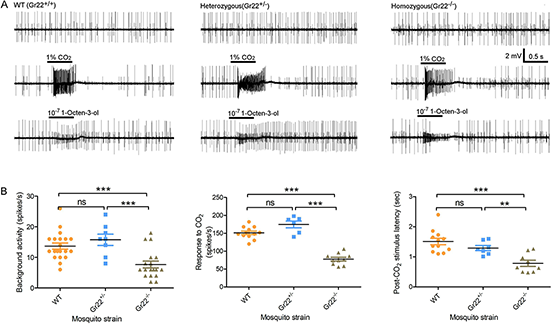
Neuronal response of wild-type, *Gr22* heterozygous and homozygous mutant mosquitoes when challenged with CO_2_. **(A)** Representative recording from capitate peg sensilla in the maxillary palp of wild-type, *Gr22* heterozygous and homozygous mutant mosquitoes; Top to bottom: background activity, response to 1% CO_2_ and response to 10^−7^ (v/v in paraffin oil) 1-Octen-3-ol. **(B)** Statistical analysis of background activity, firing frequency of cpA neuron in response to CO_2_ challenge, and latency of cpA neuron after CO_2_ challenge. Nonparametric Mann-Whitney test is applied in the statistical analysis with P<0.05 (*), P<0.01 (**) and P<0.001 (***) as significant difference.

Exploring this further, *Gr22*^*-/-*^ homozygous mutants displayed consistently deficient CO_2_ responses compared to wild-type mosquitoes at serial CO_2_ dilutions covering 3 orders of magnitude (Supplemental Figure S2A). Additionally, when exploring the temporal dynamics of CO_2_-evoked cpA neuron firing it is clear that *Gr22*^*-/-*^ mutants respond more phasically resulting in a firing pattern with a kurtosis value of 5.685 while wild-type female maxillary palp cpA neurons respond to CO_2_ more tonically giving rise to a firing pattern with a kurtosis value of 0.016 (Supplemental Figure S2B).

### *Gr22* is required for complete detection of CO_2_-mimic compounds in the cpA neuron

While in *An. coluzzii* and other mosquitoes it is likely the maxillary palp cpA neuron’s primary role is to sense host-derived CO_2_, the cpA neuron also responds to a range of non-CO_2_ odorants (Lu et al., 2007; Coutinho-Abreu et al., 2019). Insofar as such compounds represent potential CO_2_-augments and/or mimics that could be useful to further examine the mechanistic roles of Grs 22,23, and 24 as well as opportunities for optimizing odor-baited mosquito traps, we screened the *An. coluzzii* cpA neuron for responses to an in-lab chemical library of 325 compounds (Figure 5A). To begin with, stimulation with these non-CO2 compounds revealed three classes of temporally distinct cpA neuron response patterns: 1) CO_2_-like responses were elicited by compounds which not only evoked excitatory cpA responses but also gave rise to an obvious post-stimulus latency, these were identified as potential CO_2_ “mimics”; 2) Non-CO_2_-like excitatory responses without a post-stimulus latency; 3) Suppressive responses that significantly inhibited cpA neuron firing (Figure 5B, Supplemental Figure S3). Of the 50 compounds which elicited significant cpA excitatory responses, 15 including pyridine, cyclohexanone, and 2-methyl-2-thiazole display CO_2_-like mimic responses (Figure 5C, blue bars).

**Figure 5.**
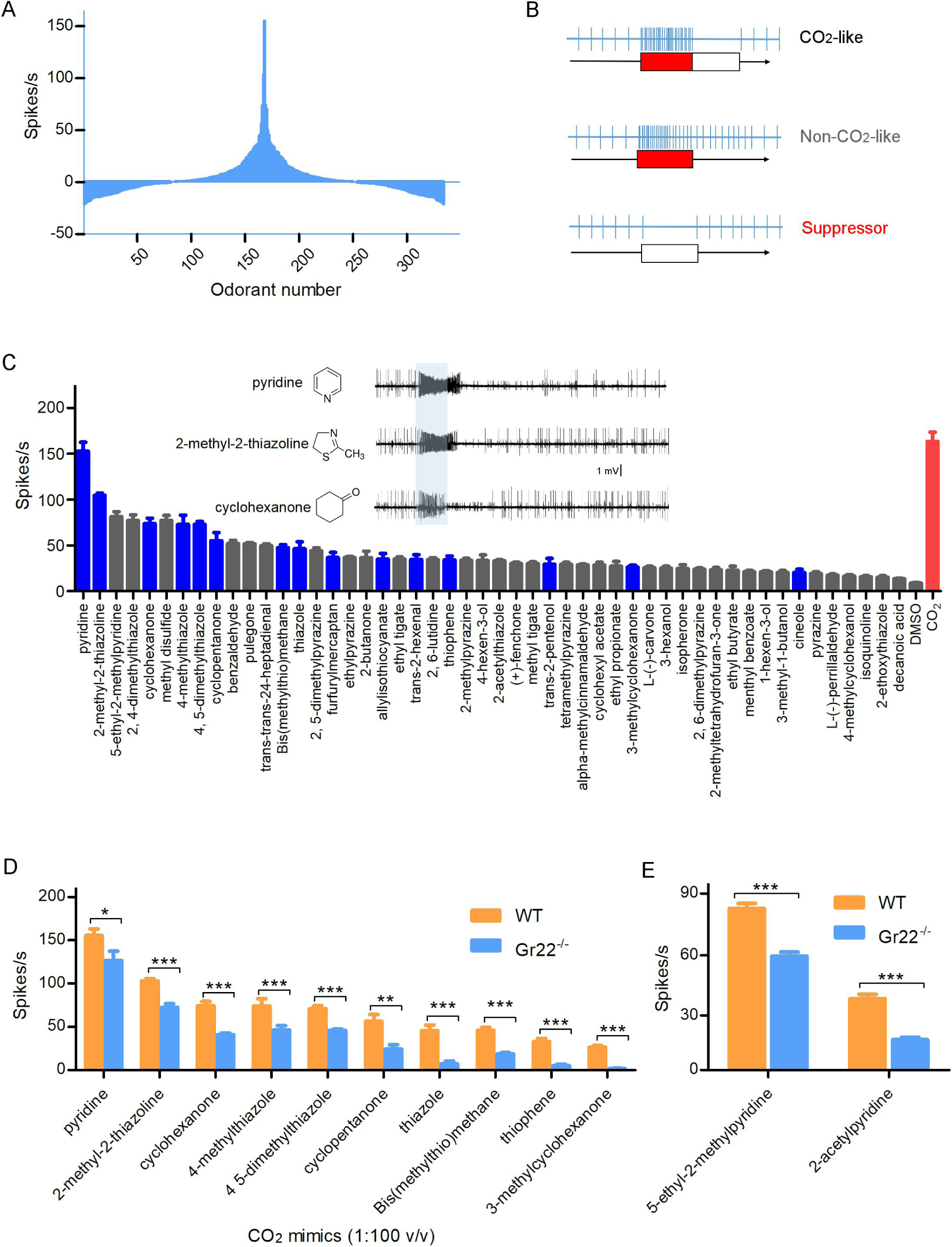
Neuronal responses of wild-type and *Gr22*^*-/-*^ mosquitoes to CO_2_-mimic compounds. **(A)** Screening the potential CO_2_-mimic compounds (10^−2^ v/v or m/v in DMSO) on the cpA neuron with our laboratory chemical library (325 compounds) and identifying the most potent compounds in activating the cpA neuron. **(B)** Classification of response pattern of cpA neuron to different compounds compared with CO_2_-evoked response. **(C)** Compounds eliciting significant excitatory response on the cpA neuron compared to the paraffin oil control. CO_2_-like responses are labeled with blue color. Non-CO_2_-like response are shown with gray color. The response to 1% CO_2_ was placed on the right end of the figure with a red bar. Top three most potent CO_2_-mimcs which elicit the strongest responses in the neuron are placed in the left corner of the figure. Representative firing signal trace of those three compounds (10^−2^ v/v) is shown. **(D)** Lower sensitivity of the cpA neuron of *Gr22* homozygous mutant mosquitoes in response to ten CO_2_ mimics compared to that in the wild-type mosquito. **(E)** Lower sensitivity of the cpA neuron of *Gr22* homozygous mutant mosquito in response to two pyridine-derived compounds compared to that in the wild-type mosquito. Nonparametric Mann-Whitney test is applied in the statistical analysis with P<0.05 (*), P<0.01 (**) and P<0.001 (***) as significant difference.

We next examined the responses of our *Gr* mutants to a panel of ten CO_2_ mimics all of which evoked strong responses in the cpA neuron of wildtype mosquitoes. As expected, the largely silent cpA neuron of *Gr23*^−/-^ and *Gr24*^−/-^ mutants were indifferent to these compounds, (Supplemental Figure S4), as well as to stimulation with compounds that evoke non-CO_2_-like excitatory responses in wild-type cpA neurons (Supplemental Figure S5). However, all ten of our CO_2_ mimics stimulated obvious, albeit significantly reduced cpA responses in *Gr22*^−/-^ mutants as compared to wild-type (Figure 5D). Of these, one heterocyclic compound, pyridine, displayed the strongest response in *Gr22*^−/-^ mutants which was nevertheless still weaker when compared to wild-type firing rates (Figure 5D). Two other pyridine-derived compounds, 2-acetypyridine and 5-ethyl-2-methylpyridine, also evoke moderate cpA spike trains in both wild type and, at lower frequencies in *Gr22*^*-/-*^ mutants (Figure 5E). These data do not seem to align with recent studies indicating that pyridine triggers stronger responses in the *Drosophila* ab1C neurons heterologously expressing *Ae. aegypti Gr2* and *Gr3* but lacking *Gr1* as compared to similar preparations expressing *Ae. aegypti Gr1, Gr2* and *Gr3* (Kumar et al., 2020).

## Discussion

We have used *in vivo* gene-targeting in *An. coluzzii* to examine the requirements of the CO_2_ receptor triad encoded by *Grs22, 23* and *24*. These genes and the putative proteins they produce are highly conserved, but interestingly not universal components of insect CO_2_ receptor complexes that underlie electrophysiological responses on the maxillary palp and other chemosensory appendages. Importantly, our data is broadly consistent with previous studies that directly place these Grs at the heart of CO_2_ responses in fruit flies and mosquitoes (Lu et al., 2007, Kwon et al., 2007, McMeniman et al., 2014). While the results from our work are, for the most part, consistent with the recent findings in *Aedes* and *Culex* (Kumar et al., 2020; Xu et al., 2020), there are several inconsistencies. To begin with the aligned data, it can now be said with confidence that in mosquitoes: (1) *Gr23* (*Gr2* in *Aedes* and *Culex*) is not a redundant triad component but rather is necessary for comprehensive responses to diverse CO_2_ stimuli and, in all likelihood, for successful reproduction; (2) *Gr23* and *Gr24* (*Gr2* and *Gr3* in *Aedes* and *Culex*, respectively) together form a partially functional complex that is able to broadly detect CO_2_; (3) the presence of *Gr22* (*Gr1* in *Aedes* and *Culex*) is required for the complete functionality of the *Gr23/24* CO_2_-sensing complex (Figure 6).

**Figure 6.**
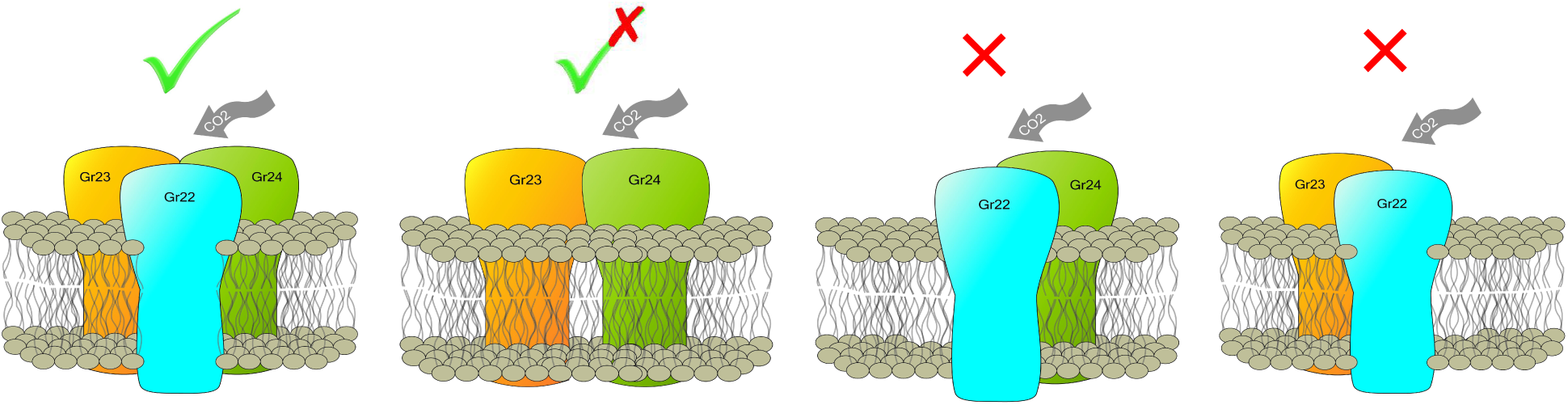
Model for organization of Grs in CO_2_ reception. While Gr23 and Gr24 are obligatory components in detecting CO_2_, Gr22 acts as an enhancer for the Gr23/24 complex in response to CO_2,_ especially at the lower concentration.

Regarding the inconsistencies, we report that *in vivo An. coluzzii Gr22*^*-/-*^ mutant maxillary palp cpA neurons display uniformly decreased sensitivity to CO_2_, CO_2_ mimics as well as to odorants that evoke non-CO_2_-like excitatory responses. In genetically engineered heterologous expression complexes utilizing the “empty” *Drosophila* ab1c neuron, lack of *Ae. aegypti Gr1* also decreases the sensitivity to CO_2_ while increasing sensitivity for mimics such as pyridine (Kumar et al., 2020). The most parsimonious explanation for this minor discrepancy is that while it is clearly faithful in many regards, *Drosophila* empty neuron systems may simply not express, traffic or otherwise utilize mosquito *Gr* genes with the same competence as observed *in vivo*. Furthermore, studies of *Culex Gr1* (Xu et al., 2020) suggest that it most likely acts as a negative modulator of the Gr2/3 complex, which directly contradicts the *Drosophila*-based studies of *Aedes* Gr1 as well as our *in vivo* findings in *Anopheles* that demonstrate that Gr22 enhances the sensitivity of the Gr23/24 complex to CO_2_. Once again, this is most likely due to the limitations of the *in vitro Xenopus* expression system and underscores the relevance of *in vivo* gene-targeting studies.

The first mosquito with a CO_2_-sensitive *Gr* gene knockout was generated in *Ae. aegypti Gr3*, the ortholog of *Anopheles Gr24* (McMeniman et al., 2014). As we now report for *Gr24*^−/-^ mutants in *An. coluzzii*, the cpA neuron in Gr3-deficient *Ae. aegypti* was shown to be completely silenced exhibiting neither background activity nor any response to CO_2_ or other compounds that typically evoke excitatory responses. This same phenomenon was observed in two studies using *Drosophila* expression platforms that lacked the fly CO_2_ receptor gene *Gr63a*, which is orthologous to *Gr24* in *Anopheles* (Jones et al., 2007; Kwon et al., 2007). Taken together, these studies across broadly diverged dipteran insects confirm the absolute functional requirement of the *Gr63a/Gr3/Gr24* component of the CO_2_-sensing apparatus.

As was the case for *Gr24*^−/-^, maxillary palp cpA neurons are completely silent in *An. coluzzii* mutants lacking *Gr23* demonstrating that loss of either gene is sufficient to render these neurons inactive across a wide spectrum of normally excitatory stimuli. Taken together, these data demonstrate that in *Anopheles*, and by extension many other insects, CO_2_-sensitive receptor complexes require both Gr23 and Gr24 as complexes lacking either receptor are inactive. This suggests that Gr23 and its homologs which are notably absent in the fully functional *D. melanogaster* CO_2_ receptor complex nevertheless provides an essential functionality to these divergent receptors. In light of the narrow CO_2_ specialization of the *Drosophila* Gr21a/63a complex (Kwon et al., 2007) it is reasonable to suggest that *Gr2/23*’s role in the mosquito receptor is related to its broader sensitivity to a wide range of non-CO_2_ stimul i(Lu et al., 2007, Coutinho-Abreu et al., 2019). While we have been unable to uncover any cpA neuron electro-physiological distinctions between the equally inactive *Gr24*^−/-^ and *Gr23*^−/-^ mutants, the unique inability of *Gr23*^−/-^ mutants to self-reproduce in laboratory mating studies (data not shown) further indicates there are nevertheless important functional distinctions between these complexes that may be developmental or otherwise salient for mosquito reproduction.

Our data suggests that in *An. coluzzii*, loss of Gr22 alters both the firing intensity and temporal dynamics of the cpA neuron’s response to CO_2_ stimulation characteristics that have been implicated in the physiological coding of compounds in the peripheral neuronal system of insects (De Bruyne et al., 2001; Hallem & Carlson, 2004). Indeed, changes in either firing frequency or temporal dynamics have been shown to influence the secondary processing in the antennal lobe as well as decision-making in the mushroom body of insect brains, which ultimately impair normal odorant perception in insects (Owald et al., 2015; Silbering et al., 2008; Turner et al., 2008). Therefore, while *An. coluzzii Gr22*^−/-^ mutants still respond to CO_2_, their diminished firing frequency and altered temporal dynamics are likely to have dramatic impacts on their behavioral responses to CO_2_ and their host-seeking ability.

## Conclusions

The malaria mosquito *An. coluzzii* uses three putative gustatory receptors, encoded by *Gr22, 23 and 24* to generate a triad complex to detect CO_2_ from blood meal hosts as well as environmental sources (Figure 6). Gene knockout studies now demonstrate that in the maxillary palp CO_2_-sensitive cpA neuron of *Anopheles* and other mosquitoes, *Gr22* functions as a modulatory component that enhances the sensitivity, discrimination and functionality of the obligatory *Gr23/Gr24* complex.

## Figure Legends

**Table S1:**
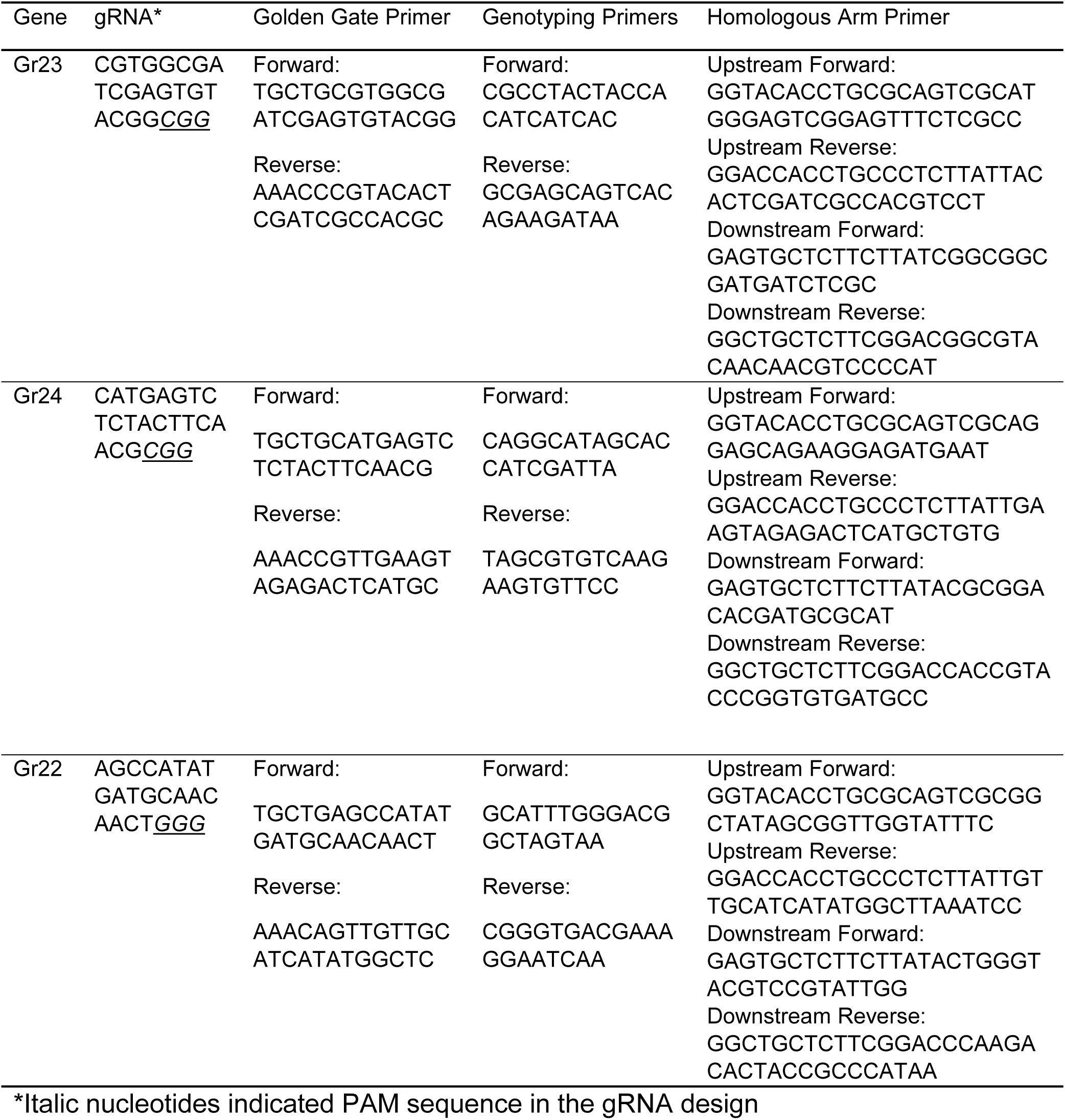
Oligonucleotides used for generating the Gr gene mutated mosquitoes

**Supplemental Figure S1.**
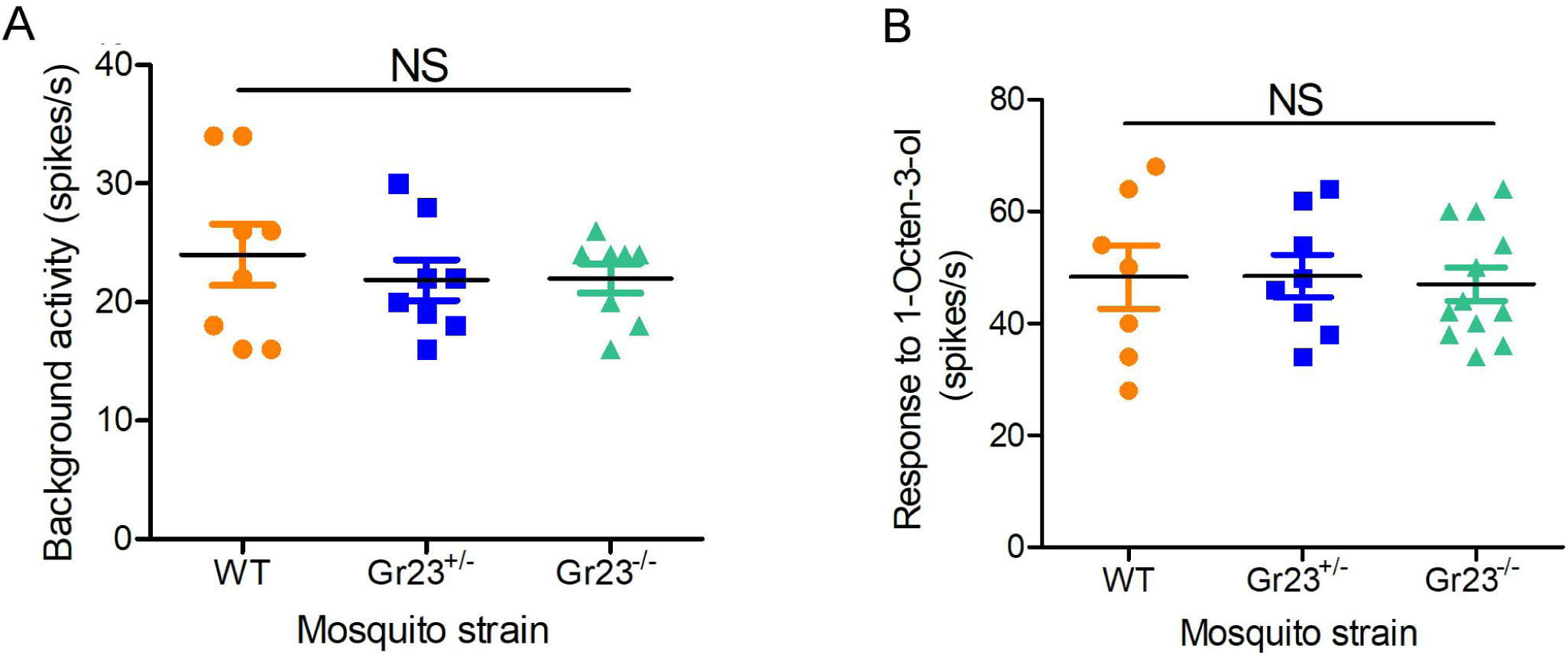
Comparison of responses of cpB/C neuron to 1-Octen-3-ol among wildtype and Gr23 mutant mosquito. **(A)** Spontaneous activity of cpB/C neuron in wildtype, *Gr23*^+/-^ and *Gr23*^*-/-*^ mosquito line. **(B)** Firing frequency of cpB/C neuron in wildtype, *Gr23*^+/-^ and *Gr23*^−/-^ mosquito line in response to 1-Octen-3-ol (10^−7^ v/v). Nonparametric Mann-Whitney test is applied in the statistical analysis with P<0.05 as significant difference, NS indicating non-significance.

**Supplemental Figure S2.**
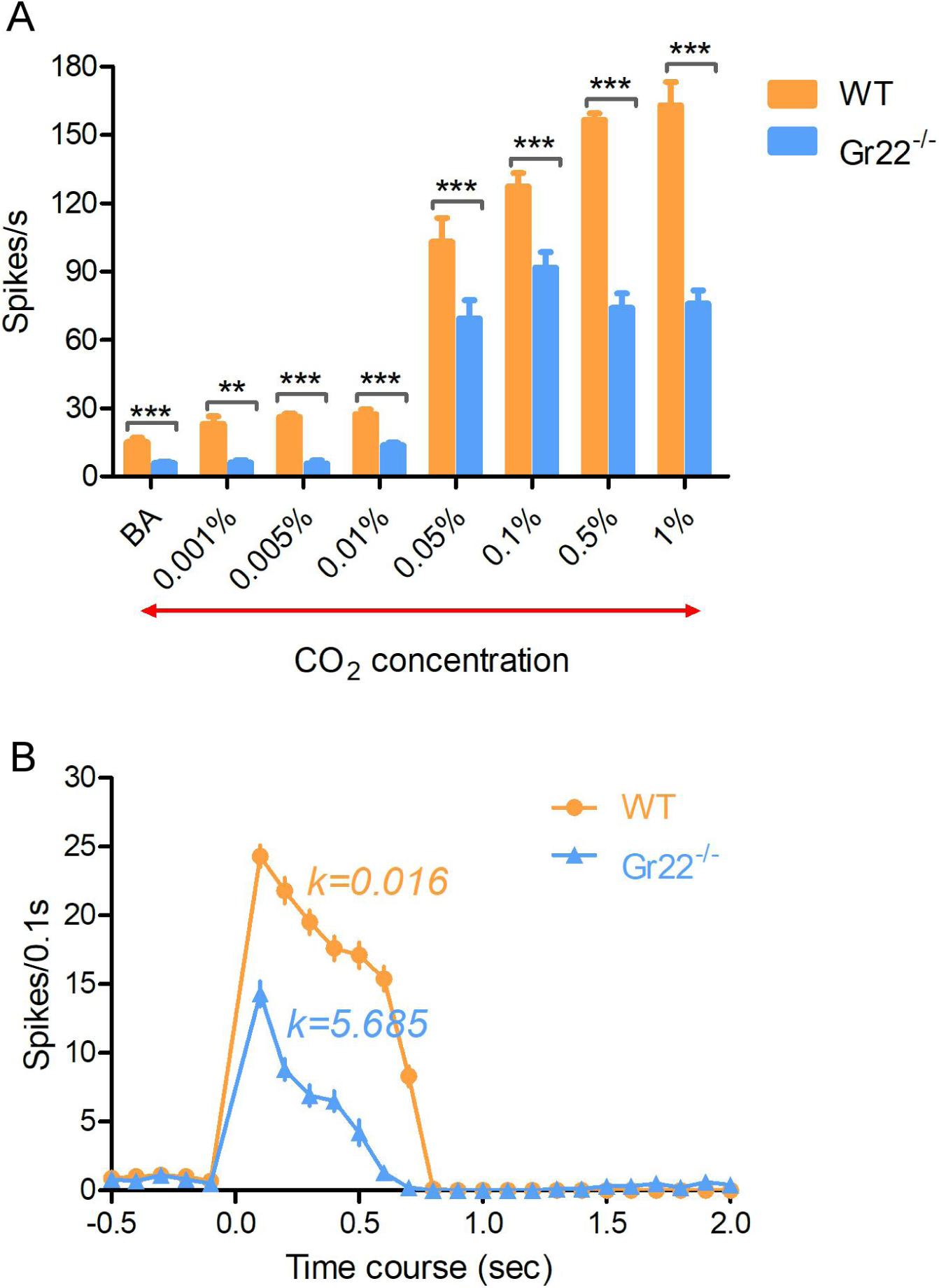
**(A)** Comparison of cpA neuron’s response to different concentrations of CO_2_ between wild-type and homozygous *Gr22*^*-/-*^ mutant mosquitoes (n=6-10). BA=background activity. Nonparametric Mann-Whitney test is applied in the statistical analysis with P<0.05 (*), P<0.01 (**) and P<0.001 (***) as significant difference. **(B)** Temporal analysis of cpA neuron’s response to 1% CO_2_ over 2 sec after stimulation. The Kurtosis value is calculated to indicate the response pattern tendency (tonic or phasic).

**Supplemental Figure S3.**
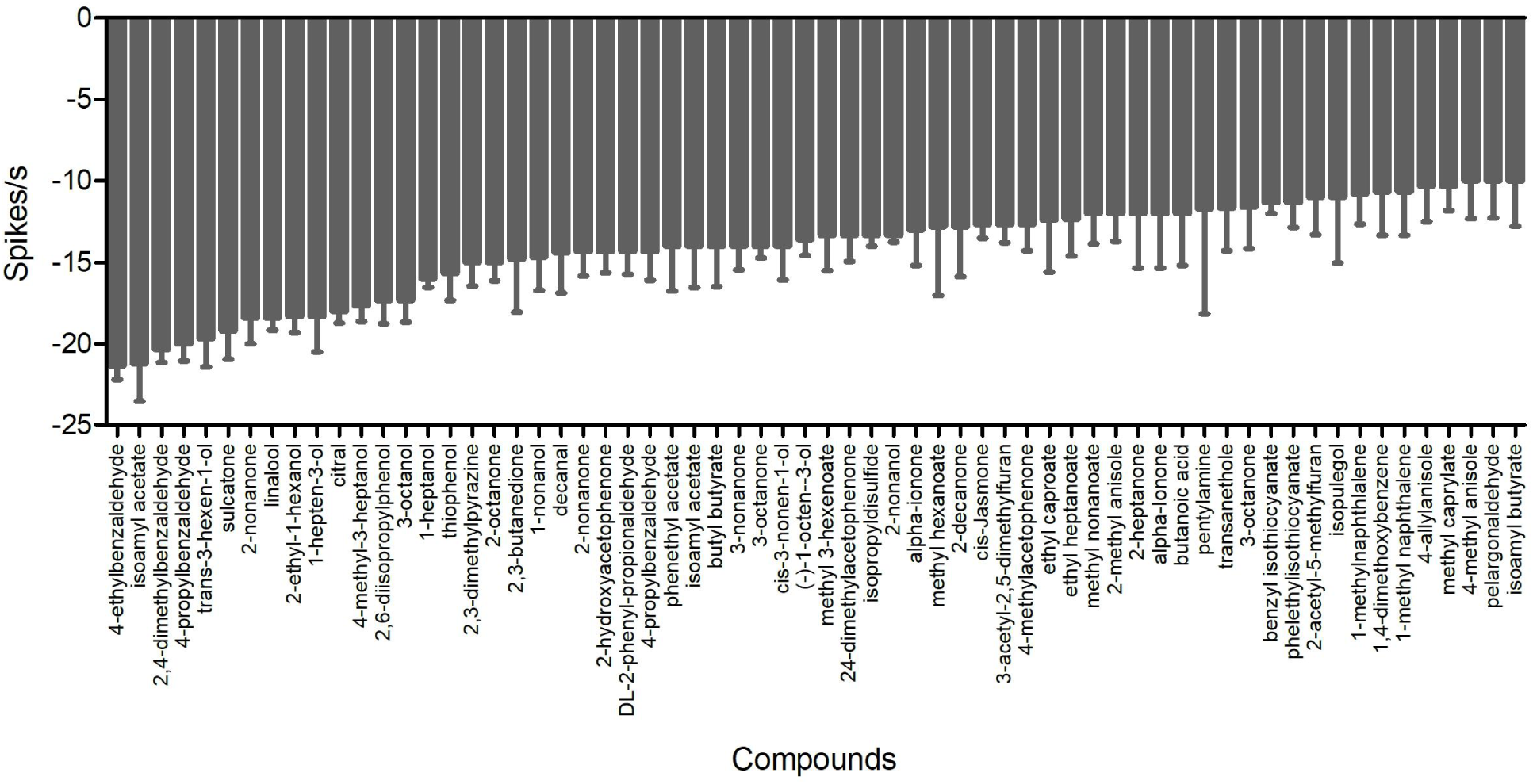
Compounds that elicited inhibitory responses on the cpA neuron with a firing frequency <=10 spikes/s (n=6-8).

**Supplemental Figure S4.**
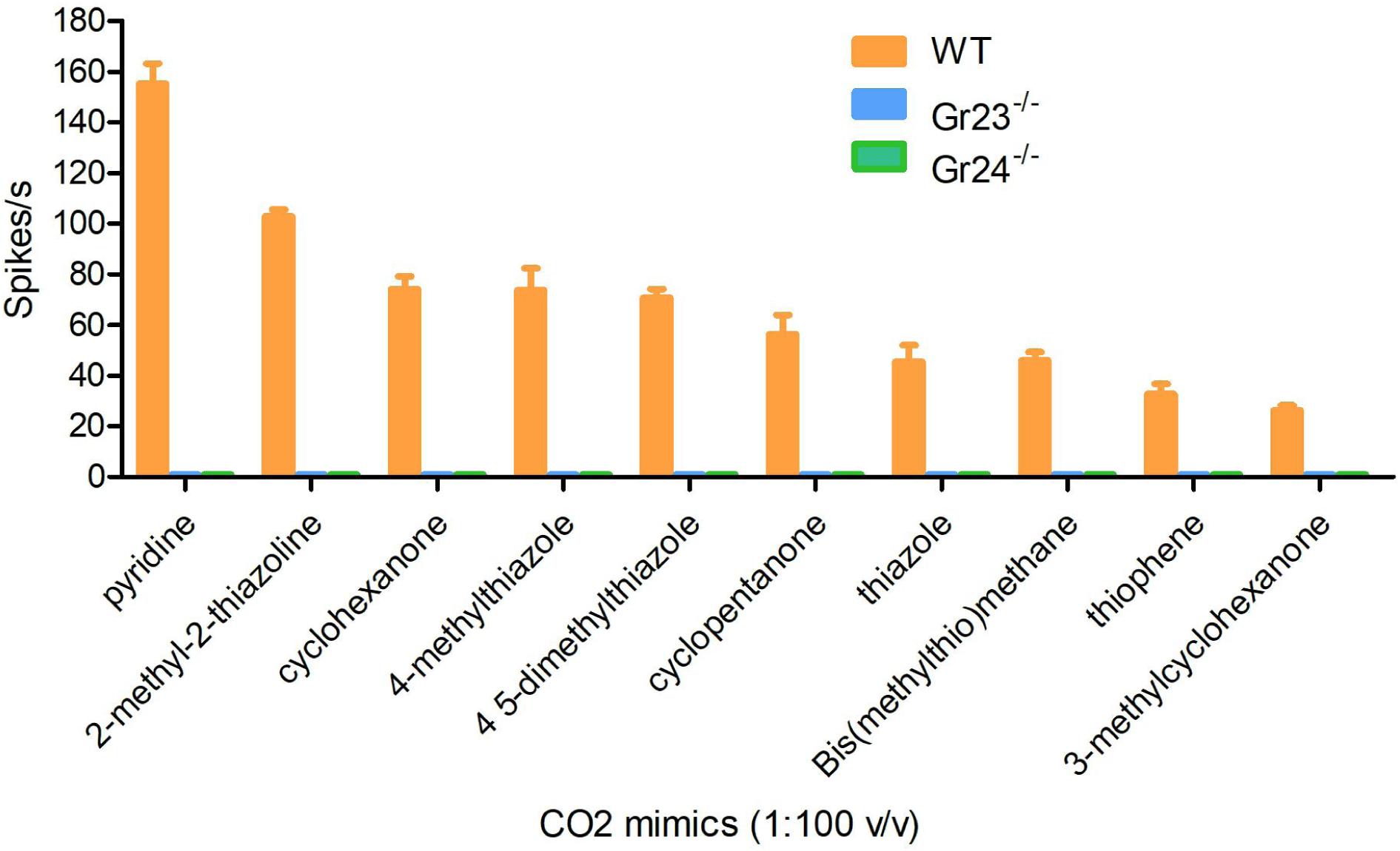
The cpA neuron of *Gr23*^*-/-*^ (n=4) and *Gr24*^*-/-*^ (n=6) mosquito presented no response to a panel of 10 CO_2_ mimics, which evoked obvious excitatory responses in that of wildtype mosquito (n=6).

**Supplemental Figure S5.**
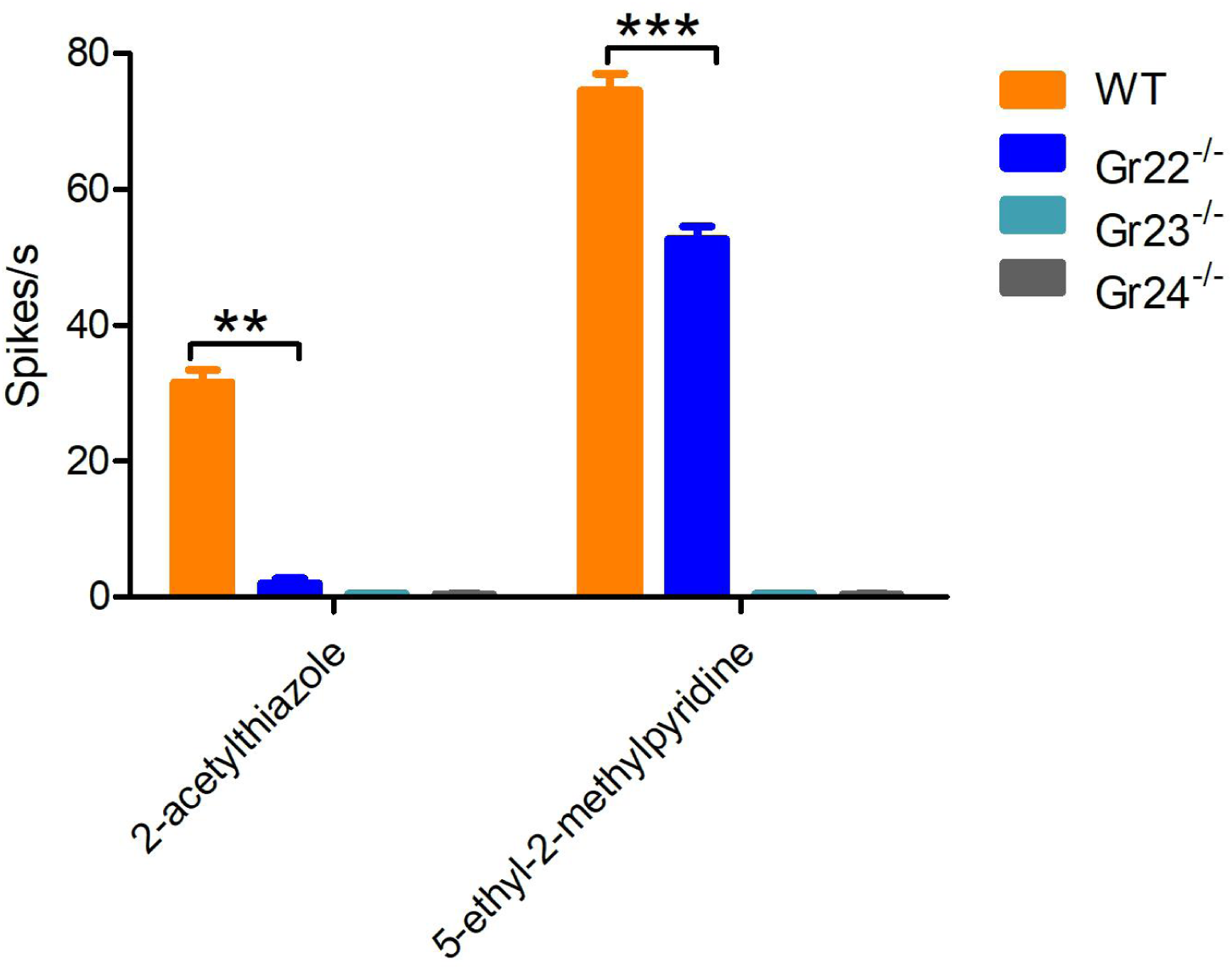
The responses of cpA neuron in *Gr22*^*-/-*^ (n=9), *Gr23*^*-/-*^ (n=4) and *Gr24*^*-/-*^ (n=6) mosquito to two non-CO2 like compounds, which evoked obvious excitatory responses in that of wildtype mosquito (n=6)

## Acknowledgments

We thank the laboratory of Dr. Andrea Crisanti (Imperial College, UK) for their generous advice for *Anopheles* gene-targeting, Dr. H. Willi Honegger, Dr. Ann Carr, Stephen Ferguson and other members of the Zwiebel lab for critical suggestions during the course of this work and acknowledge the initial studies of Dr. Pingxi Xu (University of California, Davis) in framing these experiments. We also thank Zhen Li and Samuel Ochieng for mosquito rearing and technical help. This work was conducted with the support of Vanderbilt University and a grant from the National Institutes of Health (NIAID, AI127693) to LJZ.

## Author contributions

Conceived the experiments: F.L., Y.Z. and L.J.Z.; Performed research: F.L., Y.Z., H.H.S., A.B.; Analyzed data: F.L., Y.Z. and A.B.; Wrote the paper: F.L, Y. Z., H.H.S, A.B., and L.J.Z. Approved the final manuscript: F.L, Y.Z., H.H.S., A.B. and L.J.Z.

